# Heavy metal pollutants have additive negative effects on honey bee cognition

**DOI:** 10.1101/2020.12.11.421305

**Authors:** Coline Monchanin, Erwann Drujont, Jean-Marc Devaud, Mathieu Lihoreau, Andrew B. Barron

## Abstract

Environmental pollutants can exert sublethal deleterious effects on animals. These include disruption of cognitive functions underlying crucial behaviours. While agrochemicals have been identified as a major threat to pollinators, other compounds, such as heavy metals that are often found in complex mixtures, have largely been overlooked. Here, we assessed the impact of acute exposure to field-realistic concentrations of lead, copper, arsenic, and their combinations, on honey bee learning and memory. All treatments involving single metals slowed down appetitive learning and disrupted memory retrieval at 24 h. Importantly, combinations of these metals induced additive negative effects on both processes, suggesting common pathways of toxicity. Our results highlight the need to further assess the risks of heavy metal pollution on invertebrates and to their associated ecosystem services.

**Summary statement:** Honey bees displayed reduced learning and memory performances following acute exposure to arsenic, copper or lead. Exposure to combinations of these metals induced additive effects.

## Introduction

Metal pollution is an increasingly important concern for the maintenance of ecosystems and public health (Nriagu and Pacyna, 1988). Over the last century, the widespread use of heavy metals in domestic, industrial and agricultural applications (Bradl, 2005) has considerably elevated their concentrations in water (Mance, 1987) and terrestrial habitats (Krämer, 2010; Su et al., 2014) to potentially toxic levels.

Critical pollinators, such as honey bees, are directly exposed to metal pollutants when foraging on contaminated nectar and pollen (Perugini et al., 2011; Xun et al., 2018), and while flying through air containing suspended particles (Thimmegowda et al., 2020). Metals accumulate in the bodies of adult bees (Giglio et al., 2017) and larvae (Balestra et al., 1992) as well as in the hive products (Satta et al., 2012). For instance, concomitant bioaccumulation of arsenic (As), copper (Cu) and lead (Pb), resulting from metal production industries (Kabir et al., 2012) and mining (Khaska et al., 2018; Lee et al., 2005), is common in honey bees (Badiou-Bénéteau et al., 2013; Giglio et al., 2017; Goretti et al., 2020) and their honey (Pisani et al., 2008; Terrab et al., 2005).

The deleterious effects of metals on humans are well-known (Tchounwou et al., 2012). Neural and neuromuscular alterations, sensory impairments and many related forms of behavioural dysfunctions highlight the neurotoxic effects of As, Cu, Pb and other metals (Chen et al., 2016). Deficits in cognition and memory have been reported for As (e.g. humans: Tolins et al., 2014; mice: Tyler et al., 2018; Wu et al., 2006), Pb (e.g. mice: Anderson et al., 2016; humans: Mason et al., 2014) and Cu (e.g. mice: Lamtai et al., 2020; Pal et al., 2013; flies: Zamberlan, 2020). Recent studies showed that low doses of Pb (Monchanin et al., 2020a) and selenium (Se) (Burden et al., 2016) also impair behaviour and cognition in honey bees, suggesting a widespread impact on pollinators. However, very little attention has been given to the potential combined effects of co-exposure to different metals (Monchanin et al., 2020b).

Interactions among stressors are commonly classified as antagonistic (when the effect of one stressor reduces the effect of the other one), additive (when stressors have cumulative effects) or synergistic (when stressors together have a greater effect than the sum of their individual effects) (Folt et al., 1999). Additive effects of As, Cu and Pb have been described for humans (Lin et al., 2016), rats (Aktar et al., 2017; Mahaffey et al., 1981; Schmolke et al., 1992) and fishes (Verriopoulos and Dimas, 1988). In rats, for instance, co-exposure to Pb and As disrupted brain biogenic amine levels (Agrawal et al., 2015). In humans, it was hypothesized that combined exposure to Pb and As, or other metal pollutants, have additive or synergistic toxic responses leading to cognitive dysfunction (Karri et al., 2016). To our knowledge, two studies have addressed the impact of metallic cocktails on bee physiology. Bees simultaneously exposed to Pb, cadmium (Cd) and Cu accumulated significant levels of those metals and had lower brain concentrations of dopamine compared to unexposed bees (Nisbet et al., 2018). Cd and Cu exerted a weak synergistic effect on bee survival (Di et al., 2020). However, none of these studies investigated potential effects of combined exposure on cognition.

Here we explored the effects of an exposure to single or combinations of metals on bee learning and memory. We hypothesised that combinations of metals may have synergistic negative effects, as it has been found with pesticides (Yao et al., 2018; Zhu et al., 2017). We tested individual honey bees in a standard protocol of proboscis extension reflex (PER) conditioning following acute exposure to As, Pb and Cu or a combination of them. We tested three concentrations of As, considered the most toxic substance (ATSDR, 2019), and added one concentration of Cu or Pb (binary mixtures), or both (tertiary mixture), to reach the molarity of the As solutions, allowing us to better assess any combined effects.

## Materials and methods

### Metal solutions

Arsenic (NaAsO_2_), lead (PbCl_2_) and copper (CuCl_2_2H_2_O) were purchased from Sigma-Aldrich Ltd (Lyon, France) and diluted in 50% (w/v) sucrose solution. Control bees were fed 50% sucrose solution. We used three concentrations of As (Table 1): a low concentration (0.13 μM) corresponding to the maximal permissible value in water (0.01 mg.L^-1^) (Codex Alimentarius, 2015), a high concentration (0.67 μM) corresponding to half the maximal permissible value in irrigation water (0.1 mg.L^-1^) (Ayers and Westcot, 1994), and an intermediate concentration (0.40 μM). This range of concentrations was reported in water sampled from polluted areas (e.g. mining sites) and in honey (Table S1). For Pb and Cu, we chose 0.27 μM (0.055 mg.L^-1^of Pb and 0.017 mg.L^-1^ of Cu) so that the binary combinations (As 0.13 μM + Cu 0.27 μM or As 0.13 μM + Pb 0.27 μM) could be compared to the As intermediate concentration (0.40 μM), and the tertiary combination (As 0.13 μM + Pb 0.27 μM + Cu 0.27 μM) to the As high concentration (0.67 μM) (Table 1). These concentrations of Pb and Cu have also been reported in honey samples (Table S1). All concentrations fell within sublethal ranges for the honey bee: the LD50 of elemental As for NaAsO_2_ ranged from 0.330 to 0.540 ug/bee (Fujii, 1980), the LD50 of Cu is 72 mg.L^-1^ (Di et al., 2016) and of Pb is 345 mg.L^-1^ (Di et al., 2016).

**Table 1:** Concentrations used. Combined treatments are shown in grey.

| Treatment | Molarity (µM) | Concentration (mg.L<sup>-1</sup>) |  |  | Ingestion of 5µL (ng/bee) |  |  |
| --- | --- | --- | --- | --- | --- | --- | --- |
|  |  | As | Cu | Pb | As | Cu | Pb |
| Control | 0 | 0 | 0 | 0 | 0 | 0 | 0 |
| Low [As] | 0.13 | 0.01 | 0 | 0 | 0.05 | 0 | 0 |
| [Cu] | 0.27 | 0 | 0.02 | 0 | 0 | 0.09 | 0 |
| [Pb] | 0.27 | 0 | 0 | 0.06 | 0 | 0 | 0.28 |
| Med [As] | 0.40 | 0.03 | 0 | 0 | 0.15 | 0 | 0 |
| [As+Cu] | 0.40 | 0.01 | 0.02 | 0 | 0.05 | 0.09 | 0 |
| [As+Pb] | 0.40 | 0.01 | 0 | 0.06 | 0.05 | 0 | 0.28 |
| High [As] | 0.67 | 0.05 | 0 | 0 | 0.25 | 0 | 0 |
| [As+Cu+Pb] | 0.67 | 0.01 | 0.02 | 0.06 | 0.05 | 0.09 | 0.28 |

### Bee exposure to metals

We collected honey bees (*Apis mellifera*) returning from foraging trips at the hive entrance in mornings of August 2020. We then anesthetised the bees on ice and harnessed them in plastic tubes, secured with tape and a droplet of wax at the back of the head (Matsumoto et al., 2012). We fed them 5 μL of 50% sucrose solution (see Table 1) and left them to rest for 3 h in the incubator (temperature: 25±2°C, humidity: 60%).

### Absolute learning

Prior to conditioning, we tested all bees for PER by stimulating their antennae with 50% sucrose solution, and kept only those that displayed the reflex. We then performed olfactory absolute conditioning according to a standard protocol using an automatic stimulus delivery system (Aguiar et al., 2018). Bees had to learn to respond to an olfactory conditioned stimulus (CS, 1-nonanol, Sigma-Aldrich Ltd, Lyon, France) reinforced with 50% sucrose solution, over five conditioning trials with a ten-minute inter-trial interval. Each trial (37 s in total) began when a bee was placed in front of the stimulus delivery system, which released a continuous flow of clean air (3,300 mL.min^-1^) to the antennae. After 15 s, the odour was introduced into the airflow for 4 s, the last second of which overlapped with sucrose presentation to the antennae using a toothpick with subsequent feeding for 4 seconds by presenting the toothpick to the proboscis. The bee remained another 15 s under the clean airflow. We recorded the presence or absence (1/0) of a conditioned PER in response to the odorant presentation during each conditioning trial. Bees spontaneously responding in the first conditioning trial were discarded from the analysis. The sum of conditioned responses over all trials provided an individual acquisition score (between 0 and 4), and bees responding at the last trial were categorized as learners.

### Long-term memory

After conditioning, bees were fed 15 μL of 50% sucrose solution, left overnight in the incubator, and fed another 5 μL of sucrose solution the following morning. Three hours after (24 h post-conditioning), we performed the retention test, consisting of three trials similar to conditioning except that no sucrose reward was presented. In addition to the odour used during the conditioning (CS), we presented two novel odours, in randomized order, to assess the specificity of the memory: nonanal was expected to be perceived by bees similarly to 1-nonanol, while 1-hexanol was expected to be perceived differently (Guerrieri et al., 2005). We recorded the presence or absence (1/0) of a conditioned PER to each odorant at each memory retention trial. We classified bees according to their response patterns: response to the CS only, response to the CS and the similar odour (low generalization level), response to all odours (high generalization level), no or inconsistent response.

### Statistics

We analysed the data using R Studio v.1.2.5033 (RStudio Team, 2015). Raw data are available in Dataset S1. We performed binomial generalised linear mixed-effects models (GLMM) (package lme4; Bates et al., 2015), with hive and conditioning date as random factors and treatment as fixed effect. Using the GLMMs, we evaluated whether treatment impacted the initial response to antennal stimulation, the spontaneous response in the first conditioning trial, the level of response in the last trial, the level of response to each odorant during the memory test, the proportion of bees per response pattern in the retention test, and the survival at 24 h. Note that only bees that had learnt the task were kept for the analysis of memory performance. Acquisition scores were standardised and compared with GLMMs using Template Model Builder (Brooks et al., 2017). For all response variables, we compared (1) the treated groups to the control, (2) groups exposed to concentrations of the same molarity (e.g. Med [As], [As+Cu] and [As+Pb]), (3) the separate and joint effects of the treatments (e.g. Low [As], [Cu] and [As+Cu]) in order to better identify interactive effects (antagonism, additive, synergism).

## Results and discussion

### Exposure to metals did not impact appetitive motivation

The proportion of bees that responded to the initial antennal stimulation with sucrose was similar among treatments (GLMM: p>0.05), with only 2.27% (1 out of 42 bees) of Low [As] and 4.08% (2 out of 46 bees) of [As+Cu+Pb] bees that did not respond. All bees responded in the other groups. Therefore, treatment did not affect appetitive motivation or sucrose perception, as previously reported with similar concentrations of Pb and Cu (Burden et al., 2019).

### Individual and joint exposures to metals reduced learning performances

Two out of the 381 bees submitted to the absolute learning task spontaneously responded to the first odour presentation and were therefore discarded. In all groups, the number of bees showing the conditioned response increased over trials, thus showing learning (Fig. 1A). However, fewer bees exposed to metals learned the task when compared to controls (GLMM: p<0.05), except for Low [As] (GLMM: −1.920±1.103, p=0.082). Accordingly, the acquisition scores of bees were significantly affected by treatment (Fig. 1B). Bees exposed to Med [As] (GLMM: − 0.610±0.246, p=0.013), High [As] (GLMM: −0.639±0.241, p=0.008) and [As+Cu+Pb] (GLMM: −0.592±0.244, p=0.015) had similar acquisition scores, all significantly lower than control bees. Bees exposed to [As+Pb] and Med [As] had similar acquisition scores, but bees exposed to [As+Cu] performed better (GLMM: 0.596±0.241, p=0.013). Bees exposed to High [As] and [As+Cu+Pb] exhibited similar acquisition scores (GLMM: p=0.810). We found no difference in the acquisition scores and the proportions of learners between individual and mixed treatments (GLMM: p>0.05), indicating an additive effect. Thus, exposure to metals significantly reduced learning performance, and combined exposure exerted additive deleterious effects.

**Figure 1:**
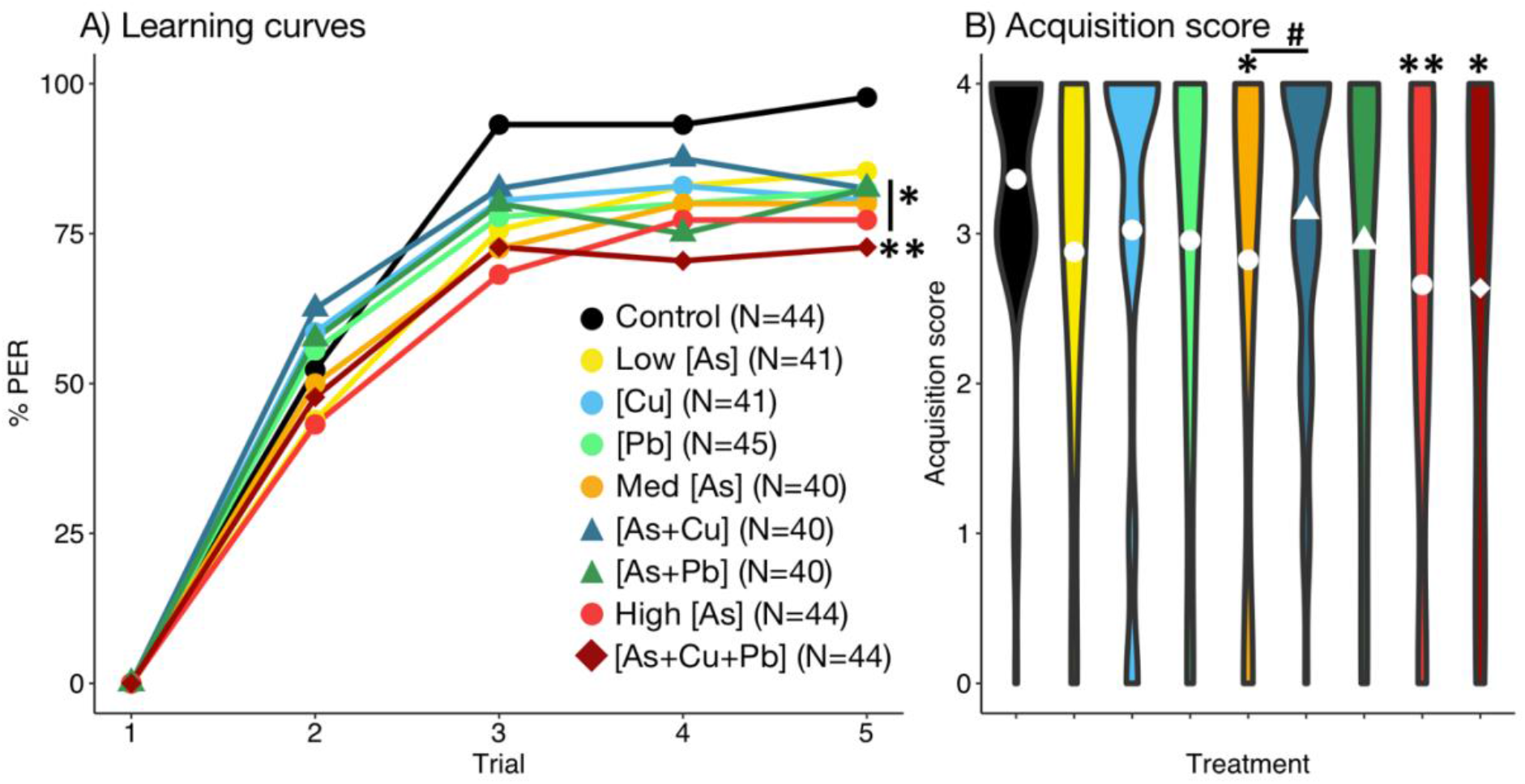
Learning. **A)** Learning curves show changes in the percentages of bees displaying the conditioned proboscis extension response (PER) over five training trials. Asterisks indicate significant differences in responses at the last trial compared to control bees. **B)** Violin plots of acquisition score values (sums of conditioned responses for each bee). Symbols (*circle*: single exposure; *triangle*: binary mixture; *diamond*: tertiary mixture) indicate the mean score for each treatment. Significant differences between groups exposed to the same molarity solutions (#) or with respect to control bees (*) are indicated (#/*p<0.05, **p<0.01; GLMM).

### Individual and joint exposures to metals reduced long-term memory specificity

We found no effect of treatment on survival at 24 h (GLMM: p>0.05). However, long-term memory was significantly affected (Fig. 2). Bees from all treatments, except Med [As] and High [As], responded less to the learnt odorant (CS) than control bees (GLMM: p<0.05) (Fig. 2A). Bees from all treatments responded similarly to the similar odour (GLMM: p>0.05). For the different odour, only bees from High [As] responded more than controls (GLMM: 1.242±0.625, p=0.047), indicating a higher degree of response generalization. We found no difference in the responses to the three different odours when comparing the groups exposed to metal solutions with the same molarity, nor between individual and mixed treatments (GLMM: p>0.05), thus indicating an additive effect.

**Figure 2:**
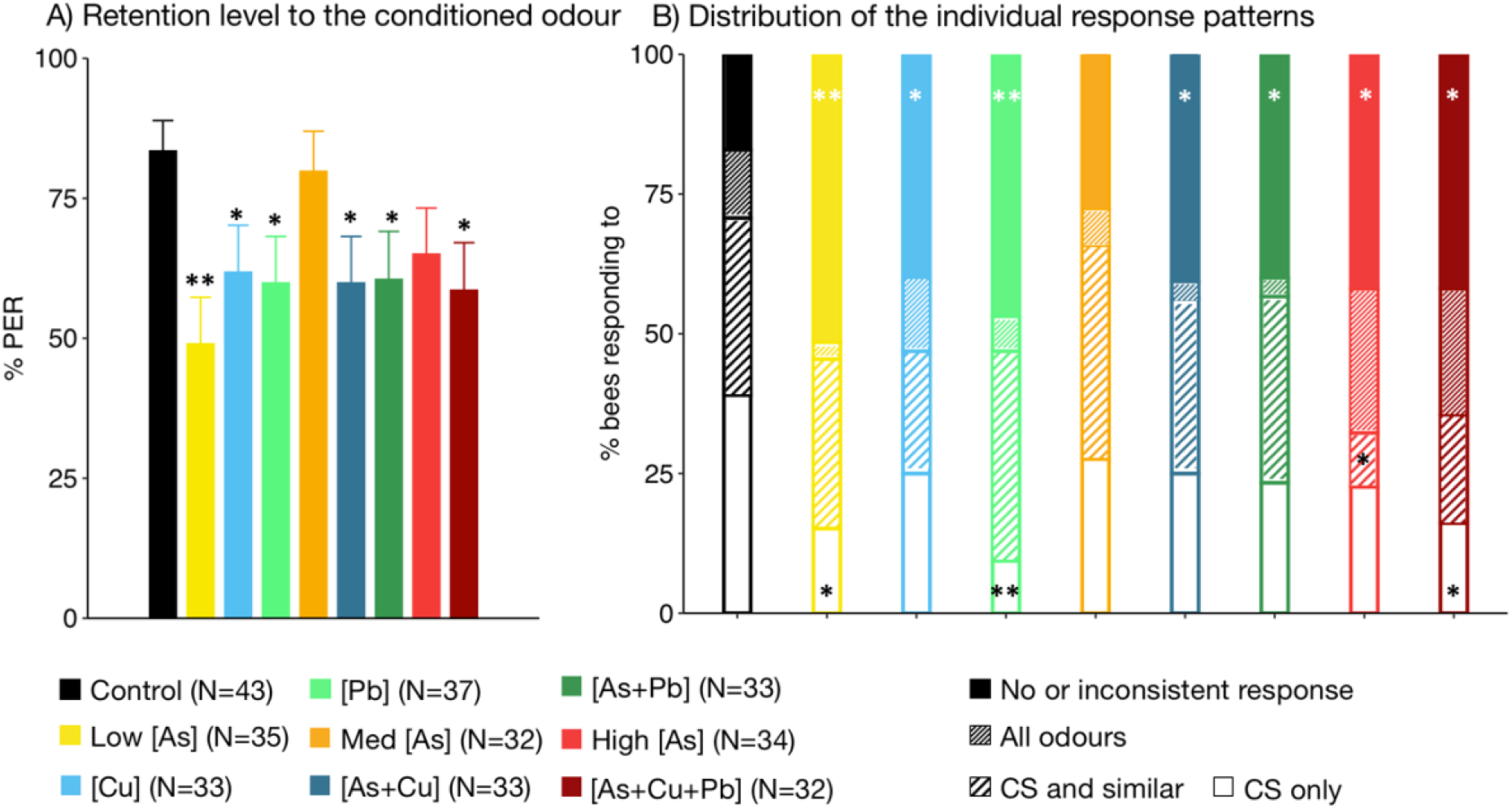
Long-term memory. **A)** Percentages of responses to the CS odour in the 24 h-memory retention test (mean ± s.e.m). **B)** Distribution of bees according to their individual response pattern during the long-term memory test: response to CS only; response to CS and similar; response to all odours; no or inconsistent response. Significant differences with controls are indicated (*p<0.05, **p<0.01; GLMM).

Individual response patterns (Fig. 2B) revealed a loss of memory specificity in bees exposed to [Pb] (GLMM: −1.795±0.690, p=0.009), Low [As] (GLMM: −1.313±0.589, p=0.026) and [As+Cu+Pb] (GLMM: −1.200±0.588, p=0.041). The distributions of individual response patterns also revealed additive effects as they did not differ among groups exposed to solutions with the same molarity, nor between single and mixed metal treatments (GLMM: p>0.05). Importantly, bees from all treatment groups but one (Med [As]) failed to respond consistently more often than controls (GLMM: p<0.05). Thus, most treatments reduced memory performance at 24h.

### The additive effects of metal mixtures may be explained by common pathways of toxicity

Although many mechanisms of metal toxicity have not yet been elucidated, some general common points already emerge from the literature. By mimicking other ions (Bridges and Zalups, 2005), metals can disrupt signalling and calcium homeostasis (particularly important in neurons) by interfering with the calcium channels (Bridges and Zalups, 2005; Chavez-Crooker et al., 2001; Tamano and Takeda, 2011). This might lead to dysfunction and cytotoxicity due to the disruption of cell signalling and calcium homeostasis. Genotoxicity (Doğanlar et al., 2014) may be achieved through covalent binding to DNA (Brocato and Costa, 2013; Senut et al., 2014). Eventually, oxidative stress and lipid peroxidation of the cell membrane may lead to neuronal death. Based on their shared mechanisms of toxicity, such as oxidative stress (Nikolić et al., 2016; Zaman et al., 1995), apoptosis (Raes et al., 2000) and interference with neurotransmitters (Nisbet et al., 2018), the toxic effects of heavy metals mixtures may be expected to be additive.

## Conclusion

We demonstrate that arsenic, lead, copper or combinations of these metals, at levels found in the environment, slow down appetitive learning and reduce long-term memory specificity. Importantly, these metals show additive effects as we found no differences between different solutions of the same molarity, suggesting that concentration was more important than identity of any specific metal for the toxicity. Learning and memory of olfactory cues play crucial roles in the behavioural ecology of bees, for the identification of profitable resources, social interactions and the recruitment of nestmates (Farina et al., 2005; Grüter et al., 2006). Therefore, ultimately, acute exposure to metal pollutants mixtures could impair fundamental hive function and population growth, a snowball effect that may be even more critical in many solitary bees that cannot rely on new cohorts of workers to replace contaminated individuals and engage in food provisioning (Klein et al., 2017).

## Acknowledgments

We thank Olivier Fernandez for assistance with beekeeping.

## Competing interests

The authors declare no competing or financial interests.

## Funding

This work was supported by the CNRS. CM was funded by a PhD fellowship from French Ministry of Higher Education, Research and Innovation. ABB was funded by a Future Fellowship from the Australian Research Council (FT140100452) and the Eldon and Anne Foote Trust. ML was funded by grants of the Agence Nationale de la Recherche (ANR-16-CE02-0002-01, ANR-19-CE37-0024) and the European Regional Development Fund (project ECONECT).

## Data availability

Raw data will be available on Dryad repository upon publication.

